# Exploring automated Feature Selection for Model-based and Density-based clustering with application to NCI 60 data

**DOI:** 10.1101/2024.04.21.589433

**Authors:** Suruchi Jai Kumar Ahuja

## Abstract

A major objective of clustering is to identify groups in the data that maximizes the similarity between objects within the same cluster and minimizes the similarity between different clusters. A challenge for data clustering, and unsupervised learning in general, is that there is often no mechanism for feature selection. In contrast, supervised learning problems can be solved in connection with feature selection methods, such as subset selection or LASSO like penalties. However, variable selection in unsupervised learning problems is not well defined since there is no response variable, which makes subset selection is far more challenging.

Consequently, there have been comparatively few methods that automate feature selection for clustering. Typically, when faced with high dimensionality, or the possibility of irrelevant features, an investigator will employ dimension reduction techniques with standard clustering algorithms. In this work, I examine two methods that encode feature selection into the clustering process, cluster variable selection via model-based clustering and density-based clustering, using the Clustvarsel and DBSCAN packages in the R programming language. These methods were applied to the NCI-60 data and compared to the Principal Component based k-means over different parameter settings. Results indicate major advantages in the performance of PC based k-means when compared to feature selection via Clustvarsel and DBSCAN, and major limitations in Clustvarsel.

## Introduction

Data mining is an interdisciplinary field, which is primarily concerned with the extraction of information from data for knowledge and discovery. The Data Mining methods and tools help in sorting and identifying patterns by establishing relationships and can be broadly classified into the two areas of supervised learning and unsupervised learning [14].

In general terms a group of data objects is defined as a cluster. The overall objective of a clustering method is to maximize the similarity between objects within the same cluster and minimize the similarity between objects of different clusters. A large spectrum of clustering methods exist and each has a slightly different criteria to measure similarity and rules for cluster assembly [17].

It is generally regarded as a tool for exploratory data analysis that discerns patterns in the data. Several clustering paradigms exist, for example connectivity models, centroid models, distribution models, density models, group models and graph based models [15].

Despite the large number of methods available for clustering, the universal requirement is that the input must be explicitly specified. However clustering has no mechanism for differentiating between relevant and irrelevant variables. So the choice of variables included in the cluster analysis must be underpinned by conceptual considerations. This is very important as the clusters formed can be very dependent on the variables included. In contrast, supervised learning, provides many opportunities for feature selection,e.g. subset selection and penalized regression. Even Classification and Regression Trees (CART) have inherent abilities to select subsets of important predictors. The common theme of these methods is that they are supervised problems, which have a response variable. On the contrary, in unsupervised problems,[23] there is no response variable and the subset selection is far more challenging. Comparatively, very little has been done on this area.

Feature Selection, also known as attribute or subset selection, is the automatic selection of features in the data that are most relevant to the predictive modeling problem [5]. It is a process of selecting a subset of optimal features that give rise to generalizable and accurate predictions. Feature selection serves two main purposes [1]. For example, it makes training and applying a classifier more efficient by decreasing the size of the effective feature space. Approaches may be advantageous computationally that aim for parsimony, and are easy to interpret. Secondly accuracy can be improved by eliminating redundant or irrelevant features [2].

In this project, we are focused on the unsupervised learning problem of identifying patterns in the data. Its goal is to partition the data into several groups, with an observation within each group to share common attributes. Model based clustering and density based clustering techniques will be worked upon the data set for a comparative analysis. In these settings, the irrelevant features are removed without too much loss of information. Also the results will be compared to PCA based k-means to getter better insights into the classical approaches vs feature selection approaches. Feature Selection is fit into these models to enhance generalization by reducing over fitting as it reduces the variance of the variables.

## Methods

### Model based Clustering Algorithm

Model based clustering is based on the assumption that the observed data comes from a population with several sub populations. This technique uses a method that assumes mixture model to cluster the data, which can also be used to further predict the group membership. The data is generally modeled using a standard statistical model which embodies a set of assumptions, which describes probability distributions[22]. Different statistical techniques are applied to the data, to understand the model such as the number of components, the probability distribution used to assign users to the various clusters, and the parameters of each model component. Once the model is specified, it can be used to assign each user to a cluster or to the set of clusters [10].

In this setting, a cluster is mixture of components and each component is described by a density function and has an associated probability or weight in the mixture. In finite mixture models, it is usually assumed that the variables and the functional form of mixing densities is known. In practice, the resulting probability model for clustering is often assumed to be a mixture of multivariate normal distributions, which is a generalization of the one dimensional, (univariate) normal distribution to higher dimensions [4].

Gaussian Mixture Models is a weighted sum of *G* components of Gaussian Densities [11]. There are N (refers to the sample size) p-dimensions observations *x*_1_, *x*_2_, ……..*x*_*n*_. Clusters are ellipsoidal centered at mean vector *µ*_*k*_, with other geometric features such as volume, shape, orientation that are conveyed through the covariance matrix Σ _*k*_ [4].

The multivariate Gaussian density for a cluster k is given as *f*_*k*_ (*x*_*i*_|*µ*_*k*_, Σ_*k*_) with parameters (*µ*_*k*_, Σ_*k*_) where *k* = 1, 2, 3, *G*; (where G is components of Gaussian Densities)

Here *π*_*k*_ is the prior probability that *x*_*i*_ belongs to the kth component, and

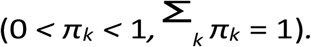

Let *a* be any constant; and then the mixture model is defined as:

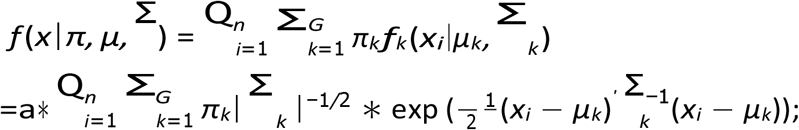

A general framework for geometric cross-cluster constraints in multivariate normal distributions was proposed [4]. The components or clusters in the mixture model have an ellipsoidal shape and are centered at *µ*_*k*_. The covariance matrices are positive symmetric, and they can be decomposed via the eigenvalue decomposition:

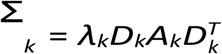

where *λ*_*k*_ is the scalar controlling the volume ellipsoidal, *A*_*k*_ is the diagonal matrix specifying the shape of the density contour and *D*_*k*_ is the orthogonal matrix which determines the orientation of the corresponding ellipsoidal. the *D*_*k*_ determines the orientation of principal components ofΣ_*k*_, *A*_*k*_ determines the shape of density contours, and *λ*_*k*_ determines the volume of the ellipsoid which is proportional to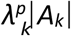 . This is the general workflow of the model based clustering algorithm.

In general, a desirable consequence of model selection is the identification of variables with more discriminating power than others. Generally the criteria based algorithms such as Hierarchical clustering algorithms are easily implemented but one of its drawbacks is that it is difficult to obtain meaningful comparison of model fit from one situation to another. In many multivariate data sets, some of the variables are redundant or irrelevant to the questions at hand, and thus do not carry much useful information about clustering [22]. The model based clustering algorithm is implemented using the clustvarsel package.

### The performance of clustering algorithms can actually be severely affected then by the presence of such variables that only serve to increase dimensionality and add redundant information. The elimination of such variables can potentially improve both estimation and clustering performance

#### The clustvarsel Package

The Clustvarsel package can be used to find the locally optimal subset of variables with cluster information in a data set with continuous variables. Clustvarsel uses a stepwise algorithm for checking the single variables for the inclusion or exclusion from the set of selected clustering variables. The method proceeds with a stepwise greedy search which checks the inclusion of each single variable not currently selected into the current set if the selected clustering variables at each step. Variables with the highest evidence of inclusion is proposed and if its clustering evidence is stronger than the evidence against clustering, it is included. At an exclusion step, it checks for the removal of each single variable in the currently selected set of clustering variables and proposes the variable with the lowest evidence for clustering. It is then removed if the evidence for its being a clustering variable is weaker than the evidence against.

The stepwise algorithm can be implemented in a forward/backward fashion, that is starting from the empty set of clustering variable and then continuing to add or remove features until there is no evidence of further clustering variables. It can also be implemented in a backward/forward fashion, that is starting from the full set of features as clustering variables and then continuing to remove or add features until there is no evidence of further clustering variables [6].

Another possible algorithm is potentially checking of less variables at each inclusion or exclusion step, and so may be quicker than the stepwise greedy search (at a possible price in terms of performance) for use on data sets with a large number of variables. At each inclusion step, this algorithm only checks single variables not currently in the set of clustering variables until the difference between the Bayesian Information Criterion for clustering versus not clustering is above a pre-specified upper level (a default value of 0 implies that the evidence for clustering is greater than that for not clustering by any amount). This algorithm is known as the headlong algorithm [8].

The Bayesian Information Criterion(BIC), is a criterion for model selection among a finite set of models; the model with the lowest BIC is preferred [7]. The evidence for a variable being useful for clustering, among the currently selected clustering variables is given by the difference in the BIC for the model based on clustering variables. The variable being checked is conditionally independent of the clustering given the other clustering variables is modeled as a regression of the variable being checked on the other clustering variables [20].

#### Density Based Clustering Algorithm

Majority of the clustering methods seek a partitioning of the data set into a predefined number of k groups where the sum of squared pairwise dissimilarities between all cluster objects with respect to a cluster representative such as a mean value, are minimized and the respective values between the clusters were maximized. These assumptions typically result in convex or spherical clusters. From a statistical point of view, theses methods correspond to parametric approaches where the unknown density of the data is assumed to be a mixture of densities, each corresponding to one of the groups in the data. Whereas, the density based clustering algorithm is very different from the parametric approaches. The density based clustering is a non-parametric approach where the clusters are considered to be areas of high density. Density based clustering algorithms do not require the number of clusters as input parameters, and there are no assumptions concerning the underlying density or the variance within the clusters [16]. As a result, density based clusters are not necessarily groups of points with a low pairwise within-cluster dissimilarity, but they have arbitrary shape [19].

Density based clusters are a set of data objects spread in the data space over an adjacent region of high density objects, separating from other density based clusters by bordering regions of low density of objects. Density based clusters can be considered as a set of points, resulting from an estimation of the probability density function for the data at a certain density level. If the level is too high, clusters exhibiting the lower density will be lost and if the level is too low, different clusters will be merged to form a single cluster [9]. The Density based clustering is implemented for feature selection using the DBSCAN Package.

### The Density Based Spatial Clustering Applications with Noise (DBSCAN) Package

In this project, a type of density based clustering, known as DBSCAN (Density based clustering applications with noise). It is a fast reimplementation of several density based algorithms of the DBSCAN family for spatial data. Spatial Data are point objects or spatially extended objects in a 2D or 3D space or in some high dimensional space [12]. Spatial data clustering is one of the most important data mining techniques for extracting knowledge from large amounts of spatial data collected in various applications. DBSCAN (Density based Spatial Clustering Applications with noise) is one of them, which discovers clusters of any arbitrary shape and can handle the noise points effectively. A drawback of DBSCAN is that it requires large volumes of memory support as it operates on the entire database. When the DBSCAN was applied to the entire NCI data set, the clusters formed weren’t very accurate and had a lot of noise points. Then Principal Component Analysis was performed on the data to reduce the dimensionality, and then the first two components were used for the DBSCAN analysis.

The DBSCAN algorithm can identify clusters in large spatial data sets by looking at the local density of database elements, using only one input parameter. Also, the user gets a suggestion on which parameter value would be suitable for the clustering. Hence, a minimal knowledge of the domain is required. DBSCAN can also determine what information should be classified as noise or outliers. The working process of the algorithm is quick and it scales very well with the size of the database, almost linearly. By using the density distribution of nodes in the database, DBSCAN can categorize these nodes into separate clusters that define the different classes.

A cluster in this model is described as a linked region that exceeds a given density threshold. The functioning of DBSCAN is directed by two well-known definitions, namely, density-reachability and density-connectability, which depend on two predefined parameter values of the DBSCAN clustering known as the size of the neighborhood denoted by *ϵ* and the minimum points in a cluster *N*_*min*_. In DBSCAN, a random point is chosen to begin with, and it finds all the points which are density-reachable from that point with respect to *ϵ* and *N*_*min*_. If the chosen point is a core point, then the formation of a cluster is completed with respect to *ϵ* and *N*_*min*_ [21]. If the chosen point is a border point, then there are no points that are density-reachable from that point, in case of which DBSCAN begins with an unclassified point to repeat the same process. The two parameters *ϵ* and *N*_*min*_ direct the notion of DBSCAN and decide the quality of clusters. These two parameters are used in a global way in the whole DBSCAN, i.e. the values of these parameters are stable for all the clusters. DBSCAN visits each point of the database, possibly multiple times.

Working of the **DBSCAN Algorithm**:

The computing process and working of the algorithm is based on the following six rules [3].

1. The Epsilon Neighborhood of a point - For a point to belong to a cluster it needs to have at least one other point that lies closer to it, than the distance Eps.

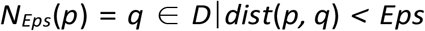
2. Directly-Density Reachable - There are two kinds of points belonging to a cluster, there are border points and core points.
3. Density-Reachable - A point is density-reachable from another point with respect to Eps and MinPts.
4. Density-connected - There are cases when two border points don’t share a specific core point. In these situations the points will not be density-reachable from each other. There must be a core point however, from which both are density-reachable.
5. Cluster - If a point x is a part of a cluster, and another point y is density-reachable from x.
6. Noise - Noise is the set of points, in the database, that doesn’t belong to any cluster.

To find a cluster, DBSCAN starts with an arbitrary point and it retrieves all points which are density reachable from that arbitrary point with respect to Eps and MinPts. If there is a core point, this procedure yields a cluster with respect to Eps and MinPts. If there is a border point then no points are density reachable from that border point and DBSCAN visits the next point of the dataset. This process is illustrated in Fig 1, which has been borrowed from [9]

**Figure 1:**
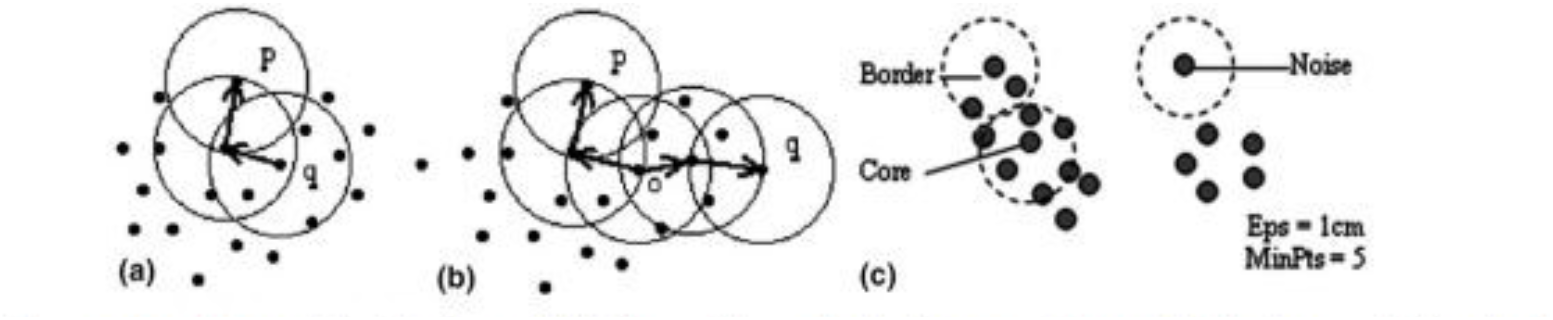
Basic Concepts of the DBSCAN Algorithm Basic concepts and terms: (a) *p* density-reachable from *q*, (b) *p* and *q* density-connected to each other by *o* and (c) border object, core object and noise.

The DBSCAN method proposed consists of computing the the k-nearest neighbor distances in a matrix of points. The algorithm calculates the average of the distances of every point to its k nearest neighbors. The value of k is specified by the user and corresponds to MinPts. The k-distances are plotted in an ascending order. The aim is to determine the knee, which corresponds to the optimal eps parameter. A knee corresponds to a threshold where a sharp change occurs along the k-distance curve. The function kNNdistplot() [in DBSCAN package] can be used to draw the k-distance plot. A k-d tree, or k-dimensional tree, is a data structure used for organizing some number of points in a space with k dimensions. It is a binary search tree with other constraints imposed on it. K-d trees are very useful for range and nearest neighbor searches. DBSCAN uses a k-d tree to find all the k-nearest neighbors in a data matrix.

### K-means Clustering

K-means clustering is one of the methods of cluster analysis in data mining. K-means clustering aims to partition n observations into k clusters in which each observation belongs to the cluster with the nearest mean, serving as a prototype of the cluster [5]. The procedure follows a simple and easy way to classify a given data set through a certain number of clusters (assume k clusters) fixed apriori. The main idea is to define k centers, one for each cluster.

The placing of these centers is very important, as different locations causes different results. The best way is to place them as far away from each other. The next step is to take each point belonging to a given data set and associate it to the nearest center. When no point is pending, the first step is completed and an early group age is done. At this point the k new centroids as centers of the clusters resulting from the previous step are recalculated. After the k new centroids are obtained, a new binding has to be done between the same data set points and the nearest new center. This procedure is iterated until covergence. The results are the k centers change their location step by step until no more changes are done or in other words centers do not move any more.

### Description of the Data - NCI Microarray Data

The US National Cancer Institute (NCI) 60 human tumor cell line anticancer drug screen (NCI 60) was developed in the late 1980s as an in vitro drug-discovery tool intended to supplant the use of transplantable animal tumors in anticancer drug screening. The screen was implemented in fully operational form in 1990 and utilizes 60 different human tumor cell lines to identify and characterize novel compounds with growth inhibition or killing of tumor cell lines. It is designed to screen up to 3,000 small molecules (synthetic or purified natural products) per year for potential anticancer activity [18].

In this project, the Human Tumor Microarray Data, also known as the NCI data from the package ElemStatLearn is used for the clustering for Feature Selection. The format of this data has 6830 columns and 64 rows, where *N* = 64 rows represents the cell lines and *P* = 6830 represents the number of genes [13]. The data set contains analysis of cell lines from 9 different cancer tissue of origin types : Breast, Central Nervous System, Colon, Leukemia, Melanoma, Non-Small Cell Lung, Ovarian, Prostate, and Renal from the NCI-60 panel. The results provide an insight into molecular mechanisms underlying the various cancer types.

The data set was processed using Principal Component Analysis, and only the first two components were used for clustering analysis. The main basis of PCA-based dimension reduction is that PCA picks up the dimensions with the largest variances. Another preprocessing step was used where a low variance filter was applied to the data, that is the variance was applied to the number of cell lines. The variances of the top 250 cell lines were used for the analysis.

## Results

The NCI data was first scaled, and then Principal component analysis was performed on the data. The first two principal components of the scaled NCI data is known as modified dataset, which was used for the K-means clustering analysis. A low variance filter was used on the NCI data and the the top 250 variances were used for the model and densitybase d clustering analysis. The resultsare shown below:

### DBSCAN

The DBSCAN algorithm was run on the processed data of the top 250 variances after applying a low variance filter. The DBSCAN algorithm was performed on the 64 gene samples. There were three clusters that were formed, which were consistent with the other findings [13]. The parameters were selected as Eps - 33.8, and MinPts - 4. The clustering contains three clusters and 5 noise points. The first cluster contained 43 tumor samples, second cluster contained 7 tumor samples, and the third cluster contained 9 tumor samples. The first cluster has more tumor samples compared to the other two clusters. The algorithm performs well as the clusters formed on the data with the low variance filter, projects well on the principal component filtered data. There are noise points that are formed are the points which do not fit well into any cluster.

The Figure 2, shows the working of DBSCAN algorithm, on the data set with a low variance filter which is plotted on the data with the first two principal components for a better visualization. There are three cluster points, each cluster assignment is represented by a different color, red green and blue. The black points represent the noise points, points which do not fit into any cluster.

**Figure 2:**
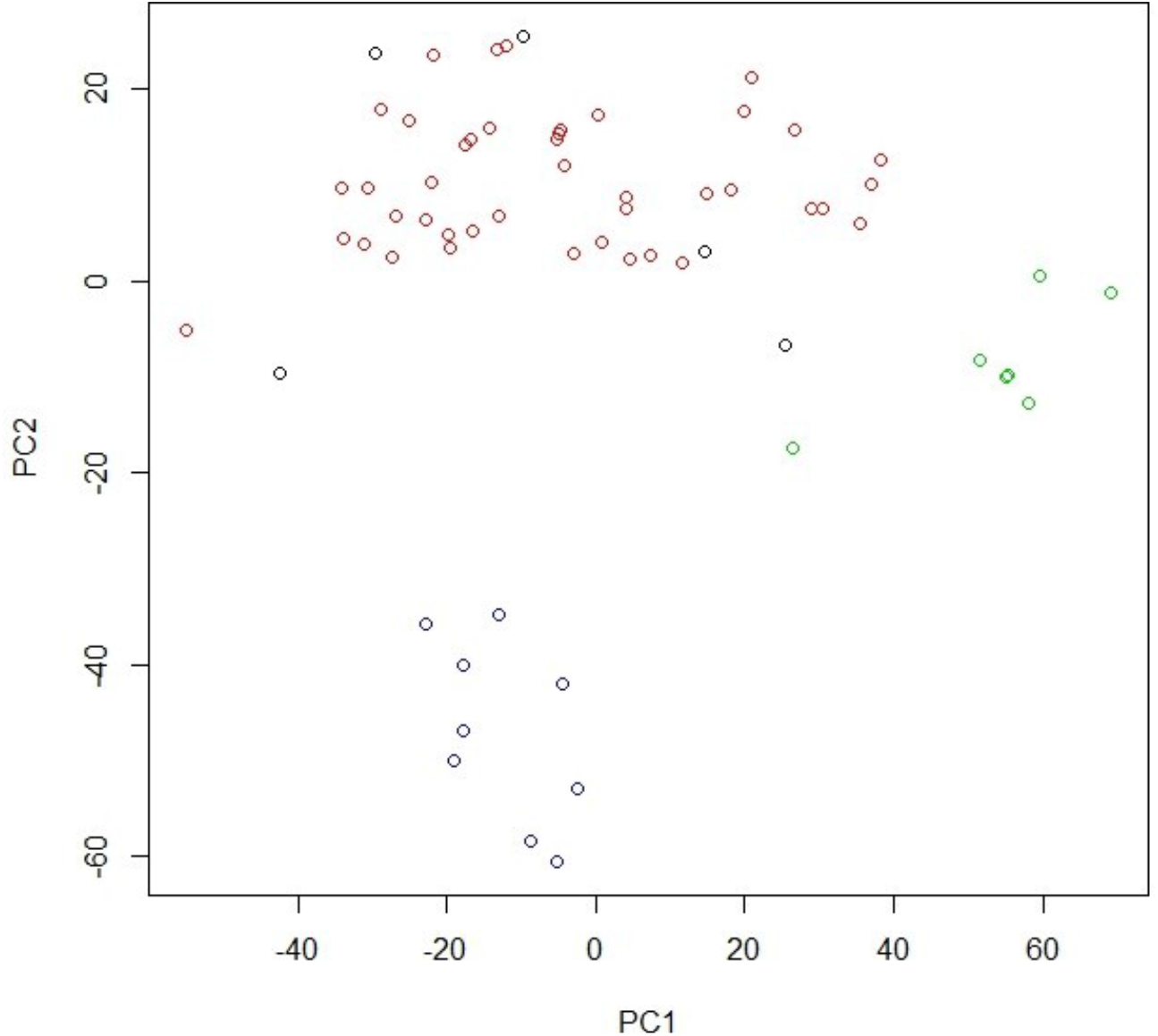
DBSCAN Algorithm on the NCI Data with a low variance filter. First two PC’s are plotted for a better visualization.

The DBSCAN algorithm uses a kd-tree, to find all the k-nearest neighbors in a data matrix (including distances) fast. This matrix of points can be used to find a suitable Eps neighborhood for DBSCAN. The k value is set to 4, which is governed by the Minpts in the options for the algorithm. This has been shown in Figure 3.

**Figure 3:**
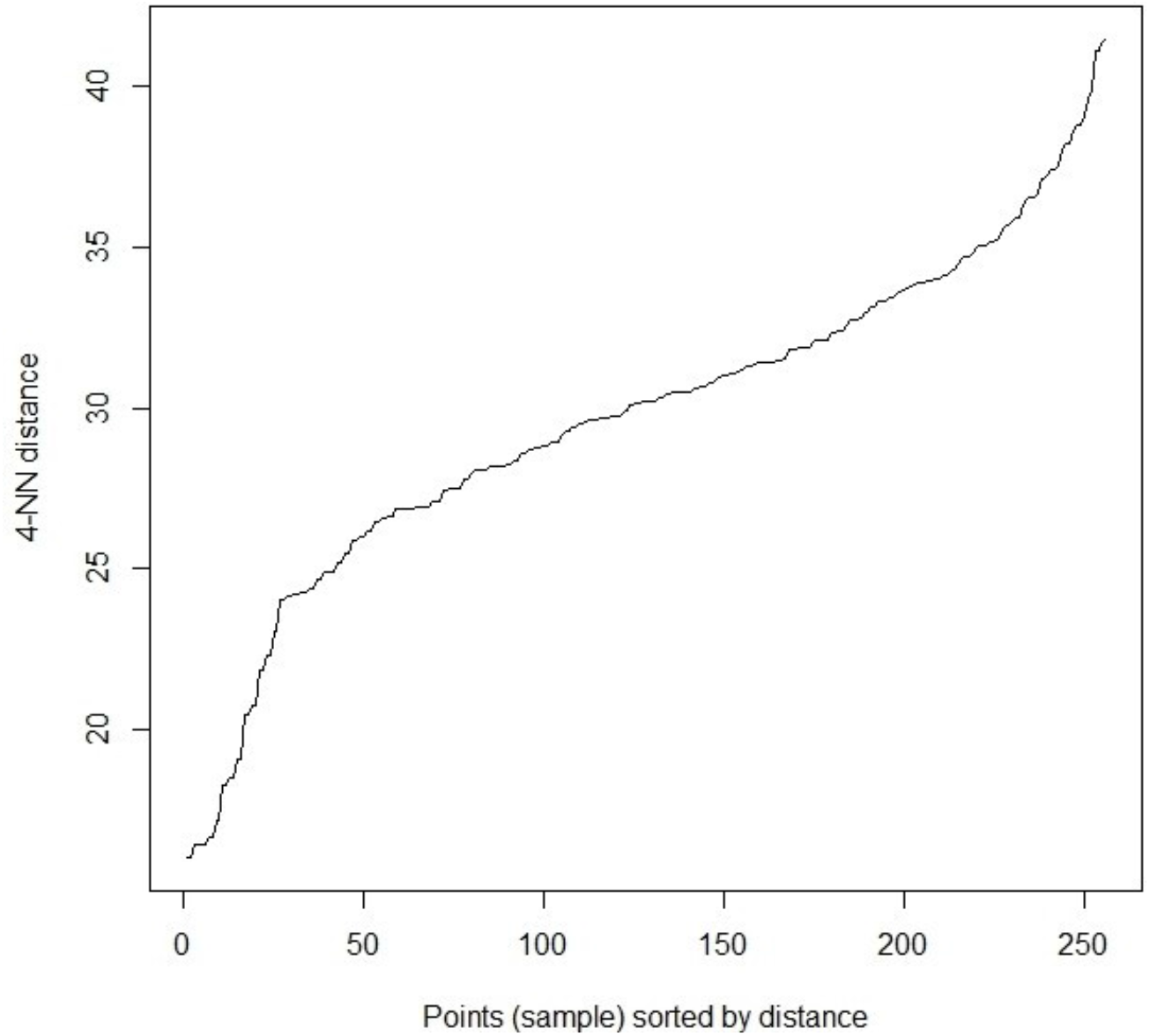
kNNdistplot on the NCI Data, calculates the k-nearest neighbor distance, and the plot helps finding a suitable value of the eps neighborhood.

A Deeper understanding of the DBSCAN Algorithm -

The results of the DBSCAN algorithm are influenced by the selection of the initial parameters which are the minimum threshold and the epsilon value. As the MinPts value increases, the number of clusters reduce and noise points increases. With different simulations, when MinPts increases to 10, there is one cluster with 36 features, second cluster with 9 features and 19 noise points, similarly when MinPts is 20, all 64 points are considered as noise points.

As the Eps value increases, the number of clusters reduce to one cluster with no noise points. As the Eps value decreases, the number of clusters increases with varying noise points. The black points in the graphs below in Figure 4, are known as noise points. Noise points are the points which don’t fit into any cluster.The Figure 4 shows how the choosing of different MinPts affects the way the clusters are formed. The red, blue and green colors represent the three clusters in each graph and the black colored points are the noise points. The last graph shows when MinPts is chosen as 20, all points are categorized as noise points.

**Figure 4:**
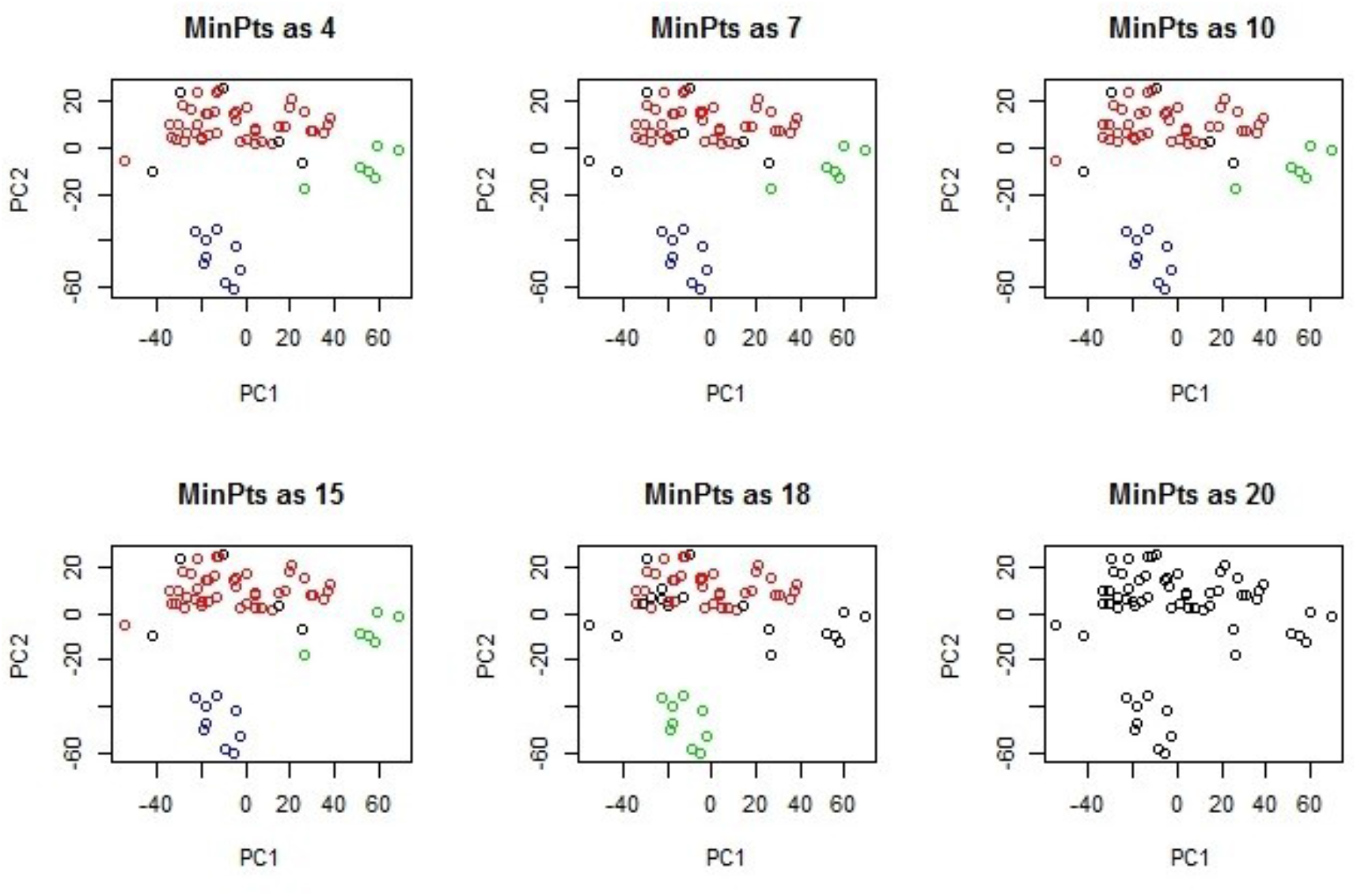
Effect of MinPts in the DBSCAN algorithm on the NCI Data, which was processed with a low variance filter.

**Figure 5:**
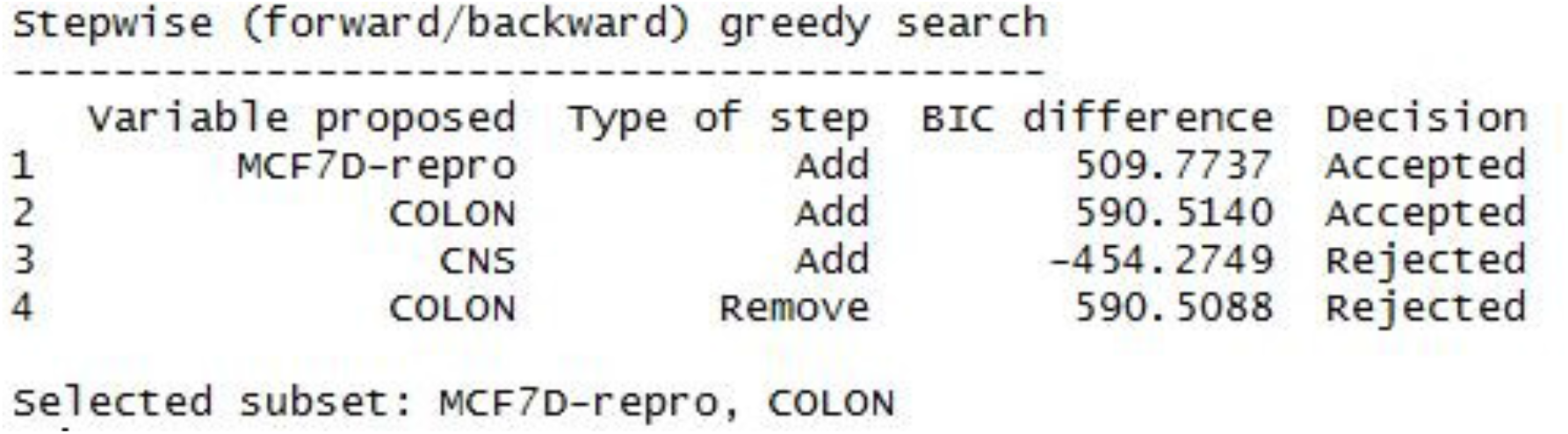
The Clustvarsel - Greedy Stepwise (Forward/Backward) Search Algorithm Output Also the same model can be achieved by using the greedy backward/forward search algorithm.

**Figure 6:**
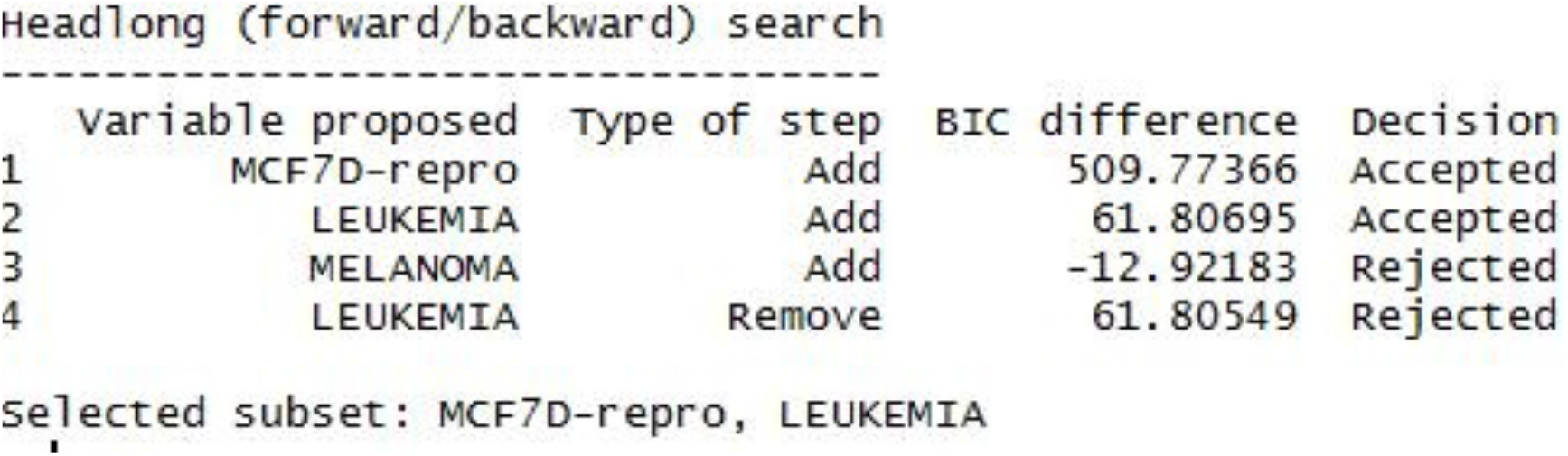
The Clustvarsel - HeadLong Search Algorithm Output

**Figure 7:**
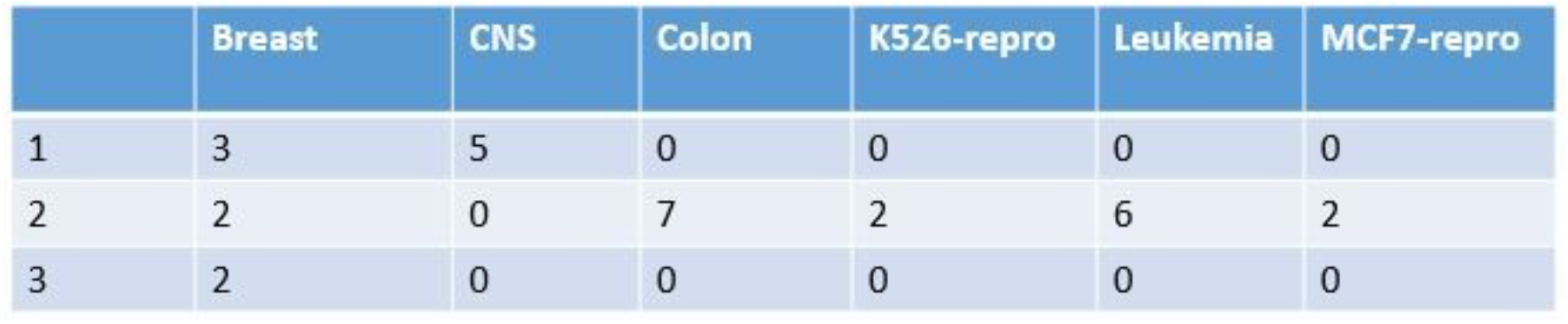
K-means Clusters

**Figure 8:**
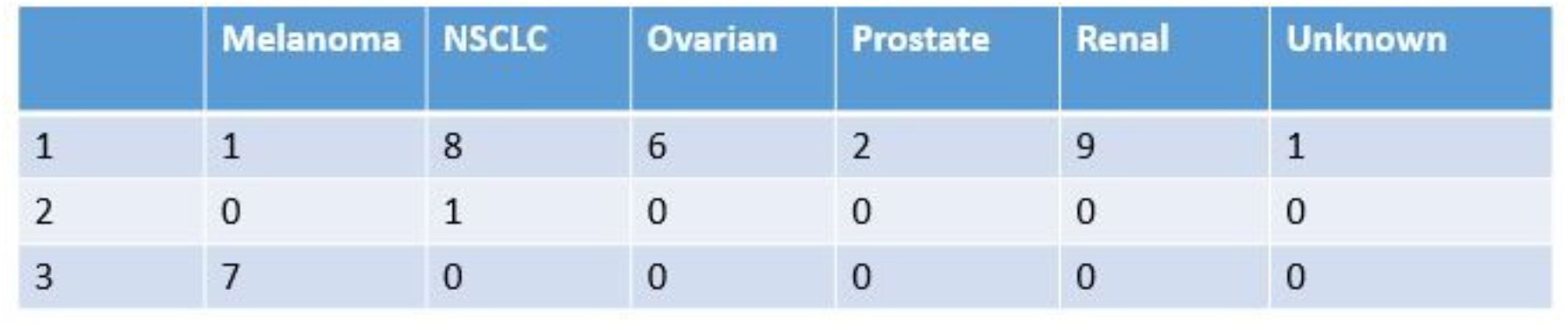
K-means Clusters-2

**Figure 9:**
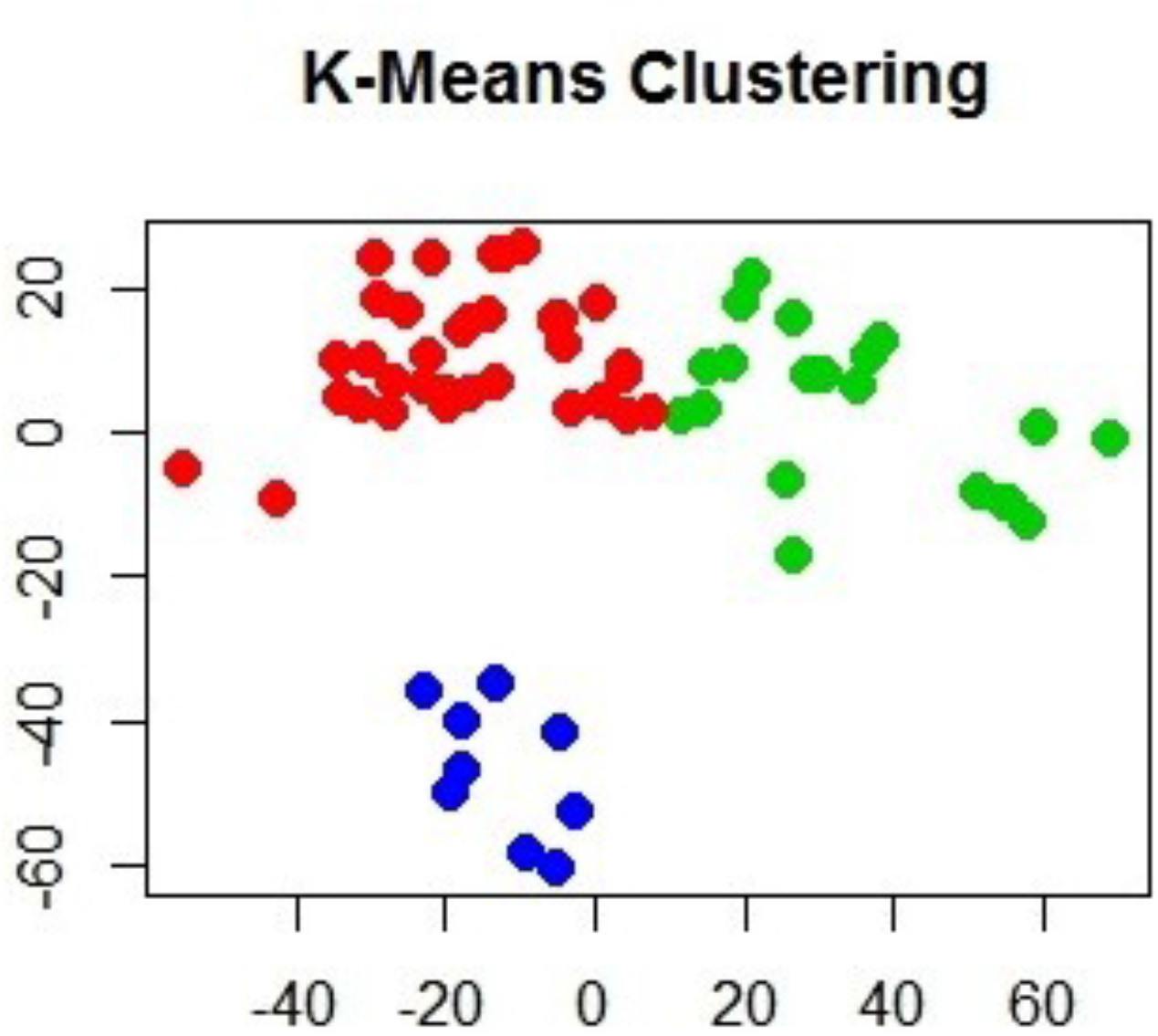
K-means Algorithm based on the PCA of the NCI Data

### Clustvarsel

The R package clustvarsel provides a convenient set of tools for variable selection in modelbased clustering using a finite mixture of Gaussian densities. Stepwise greedy search and headlong algorithm are implemented in order to find the (locally) optimal subset of variables with cluster information. The computational burden of such algorithms can be decreased by some ad hoc modifications in the algorithms, or via the use of parallel computation as implemented in the package. Also when the number of observations is large, we may allow clustvarsel to use only a subset of the observations at the model-based hierarchical stage of clustering, to speed up the algorithm.

The clustvarsel Package works with the mclust package. The Mclust model builds a Gaussian Finite Mixture Model by the EM Algorithm. Using the mclust, an ellipsoidal equal shape model with 4 components is built.

The log likelihood is -534.1785, total number of samples is 64, degrees of freedom is 17, the Bayesian Information Criterion (BIC) is -1139.058, and the ICL value is -1144.513. There are 4 clusters formed using the mclust model, where the first cluster contains 29 features, second cluster has 18 features, the third cluster has 8 features and the fourth cluster has 9 features.

Clustvarsel uses a greedy forward/backward search is used, by default. The output shows the trace of the algorithm: at each step the most important variable is considered for addition or deletion from the set of clustering variables, with each proposal which can be accepted or rejected.

The below output shows the forward/backward greedy search. There are three proposed variables : MCF7D-repro, Colon and CNS. The CNS is added into the model, but then rejected by the algorithm based on the negative BIC difference. Hence, the selected subsets are MCF7D-Rrepro and the Colon tumor samples.

### A Deeper understanding of the clustvarsel Algorithm

With data sets that contain a large number of variables, the headlong search is a much better option than the greedy search. The time taken by Headlong search is lesser when compared to the default stepwise greedy search algorithm. The Headlong search Algorithm is performed on the data, There are three proposed variables : MCF7D-repro, Leukemia and Melanoma. The Melanoma is added into the model, but then rejected by the algorithm based on the negative BIC difference. Hence, the selected subsets are MCF7D-Rrepro and the Leukemia tumor samples. The results are similar to what is achieved by the clustvarsel greedy search algorithm, as one of the selected subset is the same.

### K-means Algorithm

The k-means method has been shown to be effective in producing good clustering results for many practical applications. However, a direct algorithm of k-means method requires time proportional to the product of number of patterns and number of clusters per iteration. This is computationally very expensive especially for large datasets. K-means clustering is a method commonly used to automatically partition a data set into k groups. It proceeds by selecting k initial cluster centers and then iteratively refining them as follows:

1. Each instance is assigned to its closest cluster center.
2. Each cluster instance is updated to be the mean of its constituent instances.

The k-means algorithm was run on the new data, which was formed by performing Principal component analysis on the scaled raw data. There were three clusters that were formed. The clustering accuracy was better when performed on the PCA components of the data when compared to the entire data. The cluster observations indicate PCA dimension reduction is particularly beneficial for K-means clustering. This clustering approach is sensitive to the initial selection of centroids .

There are three clusters formed in K-means, also they are quality clusters. The first cluster has 3 subtypes of breast cancer, 5 subtypes of CNS, 1 subtype of Melanoma, 8 subtypes of NSCLC, 6 subtypes of Ovarian cancer, 2 subtypes of Prostate cancer, 9 subtypes of Renal cancer and 1 unknown cancer subtype. The second cluster has 2 subtypes of breast cancer, 7 subtypes of Colon, 2 of the K256-repro cancer, 6 subtypes of the Luekamia, 2 of the MCF7-repro cancer, 1 of the NSCLC. The third cluster has 2 of the breast cancer subtypes and 7 of the Melanoma.

Here it is not surprising to see how this clustering algorithm has captured that a few cancers are present in more than one cluster as some cancers have subtypes, and show different symptoms and features. For example, the Breast Cancer is present in three clusters, Melanoma is present in both cluster 1 and cluster 3, NSCLC is present in cluster 1 and cluster 2.

## Discussion

In DBSCAN, the border points are arbitarily assigned to clusters in the original algorithm. DBSCAN treats all the border points as noise points. Points which cannot be assigned to a cluster, will be reported as members of noise cluster. A noise point is any point that is not a border point or a core point. DBSCAN captures the insight that clusters are density based objects. Points in a cluster, their *k*^*th*^ nearest neighbors are at roughly the same distance. Noise points have the *k*^*th*^ nearest neighbors at the farthest distance.

When DBSCAN was performed on the modified data, with Eps as 33.8 and MinPts as 4, there were 5 noise points found and three clusters were formed. On visualizing the results, the cluster distribution fits well when projected on to the data with the low variance filter. The performance of this algorithm is efficient and is less time taking.

The Clustvarsel algorithm (both the stepwise greedy and headlong search) were performed on the entire data, and it takes a really long time to iterate. Due to the computational demands, we ran the clustvarsel algorithm to cluster the 64 tumor samples using the genes identified with the low variance filter. The performance of clustvarsel algorithm is terrible, as it does not provide any significant results in terms of the selected subsets. The working of the algorithm also shows how each variable is added and removed based on its BIC difference. The headlong and greedy search algorithm, were performed and different subsets were formed. This clustvarsel algorithm, takes 64 samples and reduces it to a subset of 3 variables, which is too small to be clustered. The variable selection methodolgy does not provide signifigant results to go ahead with the model based clustering, as it there not much information provided.

In conclusion, the performance of k-means algorithm is better than Density Based clustering algorithm and the Model Based clustering algorithm. The k-means algorithm forms quality clusters. The DBSCAN forms a better distribution of clusters when compared to the performance of clustvarsel. The time taken to run each clustering algorithm on the data also differs, with the k-means being the fastest followed by the DBSCAN algorithm. The Model based Clustering algorithm takes more time when compared to other two algorithms. The time factor of these clustering algorithms, also plays a very important role in the applications to show the efficiency of its working. The use of principal component analysis filter and the low variance filter on the original data, improves the efficiency and the time taken by k-means and Density based clustering algorithms.

## Notes

### Competing Interest Statement

The authors have declared no competing interest.

## References

[1] Salem Alelyani, Jiliang Tang, and Huan Liu. Feature selection for clustering: A review. Data Clustering: Algorithms and Applications, 29, 2013.

[2] Martin Azizyan, Aarti Singh, and Wei Wu. Experimental evaluation of feature selection methods for clustering.

[3] Henrik Bäcklund, Anders Hedblom, and Niklas Neijman. A density-based spatial clustering of application with noise,. Data Mining TNM033, pages 11–30, 2011.

[4] Jeffrey D Banfield and Adrian E Raftery. Model-based gaussian and non-gaussian clustering. Biometrics, pages 803–821, 1993.

[5] Christos Boutsidis, Petros Drineas, and Michael W Mahoney. Unsupervised feature selection for the k-means clustering problem. In Advances in Neural Information Processing Systems, pages 153–161, 2009.

[6] Nema Dean. The variable selection for model-based clustering (clustvarsel) package. 2009.

[7] Nema Dean and Adrian E Raftery. The clustvarsel package. R package version 0.2-4, 2006.

[8] Nema Dean and Adrian E Raftery. clustvarsel: Variable selection for model-based clustering. R package version, 1, 2009.

[9] Martin Ester, Hans-Peter Kriegel, Jörg Sander, Xiaowei Xu, et al. A density-based algorithm for discovering clusters in large spatial databases with noise. In Kdd, volume 96, pages 226–231, 1996.

[10] Chris Fraley and Adrian E Raftery. Model-based clustering, discriminant analysis, and density estimation. Journal of the American statistical Association, 97(458):611–631, 2002.

[11] Chris Fraley and Adrian E Raftery. Mclust version 3: an r package for normal mixture modeling and model-based clustering. Technical report, DTIC Document, 2006.

[12] M Hahsler. dbscan: Density based clustering of applications with noise (dbscan) and related algorithms. R package version 0.9-2, URL http://CRAN.R-project.org/package=dbscan, 2015.

[13] Maintainer Kjetil Halvorsen. Package elemstatlearn.

[14] Zhexue Huang. A fast clustering algorithm to cluster very large categorical data sets in data mining. In DMKD, page 0, 1997.

[15] Michael Kearns, Yishay Mansour, and Andrew Y Ng. An information-theoretic analysis of hard and soft assignment methods for clustering. In Learning in graphical models, pages 495–520. Springer, 1998.

[16] Hans-Peter Kriegel, Peer Kröger, Jörg Sander, and Arthur Zimek. Density-based clustering. Wiley Interdisciplinary Reviews: Data Mining and Knowledge Discovery, 1(3):231–240, 2011.

[17] William M Rand. Objective criteria for the evaluation of clustering methods. Journal of the American Statistical association, 66(336):846–850, 1971.

[18] Douglas T Ross, Uwe Scherf, Michael B Eisen, Charles M Perou, Christian Rees, Paul Spellman, Vishwanath Iyer, Stefanie S Jeffrey, Matt Van de Rijn, Mark Waltham, et al. Systematic variation in gene expression patterns in human cancer cell lines. Nature genetics, 24(3):227–235, 2000.

[19] Joerg Sander. Density-based clustering. In Encyclopedia of Machine Learning, pages 270–273. Springer, 2011.

[20] Luca Scrucca and Adrian E Raftery. clustvarsel: A package implementing variable selection for model-based clustering in r. arXiv preprint 1411.0606, 2014.

[21] Wei Wang, Jiong Yang, Richard Muntz, et al. Sting: A statistical information grid approach to spatial data mining. In VLDB, volume 97, pages 186–195, 1997.

[22] Ka Yee Yeung, Chris Fraley, Alejandro Murua, Adrian E. Raftery, and Walter L. Ruzzo. Model-based clustering and data transformations for gene expression data. Bioinformatics, 17(10):977–987, 2001.

[23] Zheng Zhao and Huan Liu. Spectral feature selection for supervised and unsupervised learning. In Proceedings of the 24th international conference on Machine learning, pages 1151–1157. ACM, 2007.

